# Mitochondrial oxidative stress drives IL-12/IL-18-induced IFN-γ production by CD4^+^ T cells and is controlled by Fas

**DOI:** 10.1101/2021.01.19.427239

**Authors:** Gorjana Rackov, Parinaz Tavakoli Zaniani, Sara Colomo del Pino, Rahman Shokri, Melchor Alvarez-Mon, Carlos Martinez-A, Dimitrios Balomenos

**Affiliations:** Department of Immunology and Oncology, Centro Nacional de Biotecnología-Consejo Superior de Investigaciones Científicas (CNB-CSIC), Madrid, Spain; Service of Internal Medicine and Immune System Diseases-Rheumatology, University Hospital Príncipe de Asturias, (CIBEREHD), Alcalá de Henares, Spain; Department of Medicine and Medical Specialties, Faculty of Medicine and Health Sciences, University of Alcalá, Alcalá de Henares, Spain

**Keywords:** mitochondrial reactive oxygen species, IFN-γ, IL-12/IL-18, Fas, effetor/memory CD4 T cells

## Abstract

Mitochondrial activation and mROS production are crucial for CD4^+^ T cell responses and have a role in naïve cell signaling after TCR activation. However, little is known about their role in recall responses driven by cytokine signaling. Here, we found that mROS are required for IL-12 plus IL-18-driven production of IFN-γ, an essential cytokine in inflammatory and autoimmune disease development. In particular, memory-like cells obtained after activation-induced differentiation showed faster and augmented mROS accumulation and increased IFN-γ production in response to IL-12 plus IL-18 compared to naïve T cells. In contrast, mROS induction was similar in naïve and memory-like cells after TCR-dependent signaling. Taken together these results suggested that memory-like CD4^+^ T cells treated by IL-12 plus IL-18 attained conditions for an extraordinary mROS-producing potential. mROS inhibition significantly downregulated the production of IFN-γ and the expression of CD44 activation marker, suggesting a direct mROS effect on the activation of memory-like T cells. Mechanistically, mROS was required for optimal activation of key signaling pathways that drive IFN-γ production after IL-12 plus IL-18 T cell stimulation, such as PKC-θ, AKT and STAT4 phosphorylation, and NF-κB activation. Notably, we identified increased mROS as key promoters of hyperactivation and IFN-γ overproduction in Fas-deficient *lpr* memory-like CD4^+^ T cells compared to WT cells, following IL-12 plus IL-18 stimulation. mROS inhibition significantly reduced the population of disease-associated CD44^hi^CD62L^lo^ *lpr* CD4^+^ T cells and their IFN-γ production. These findings uncover a previously unidentified role for Fas in regulating mitochondrial ROS production by memory-like T cells. This apoptosis-independent Fas activity might contribute to the accumulation of CD44^hi^CD62L^lo^ CD4^+^ T cells that produce increased IFN-γ levels in *lpr* mice. Overall, our findings pinpoint mROS as central regulators of TCR-independent signaling, and support mROS pharmacological targeting to control aberrant immune responses in autoimmune-like disease.

## Introduction

Upon TCR activation, mitochondria rapidly translocate to the immunological synapse, leading to electron transport chain (ETC) activation (1) and mitochondrial reactive oxygen species (mROS) production. Superoxide (O_2_^—•^) is generated at mitochondrial complexes I and III (2, 3), from where it can reach cytosol either directly through mitochondrial membrane or after conversion to hydrogen peroxide (H_2_O_2_) (4). It has been established more recently that mROS play crucial role as redox signaling molecules in T cells, regulating IL-2 and IL-4 production upon TCR triggering (5, 6).

IL-12 and IL-18 are required to boost T_H_1 responses and IFN-γ production after TCR ligation (7, 8). In a mechanism evolved to provide an early source of IFN-γ and contribute to the innate immune response, IFN-γ can be induced in a TCR-independent manner, only by IL-12 and IL-18 (9, 10). It has been proposed that this “cytokine-induced cytokine production” promotes chronic inflammation and autoimmunity (11, 12). Indeed, we previously demonstrated that IFN-γ production by *lpr* T cells creates a loop by increasing the macrophage inflammatory potential (13). The signaling mechanisms that drive IFN-γ production in response to IL-12 plus IL-18 are distinct compared to those downstream of TCR, and lead to differential NF-κB recruitment to *Ifng* locus (14–16).

Although the role of mROS has been greatly appreciated downstream of TCR, their role in recall responses of previously activated CD4^+^ T cells and in IL-12 plus IL-18-dependent signaling regulation remains unexplored. Excessive production of ROS is thought to underlie some of the pathological immune reactions implicated in autoimmunity. T cells from lupus patients exhibit mitochondrial hyperpolarization, resulting in increased mROS production (17, 18). In a previous study, we demonstrated that autoimmune traits in *lpr* mice were decreased by controlling autoreactive T cell hyperactivation and their IFN-γ overproduction (13). However, the direct link between mROS and IFN-γ hyperproduction in autoreactive T cells has not been made thus far.

Here, we hypothesized that mROS act as driving force to modulate signal transduction leading to hyperproduction of IFN-γ. We analyzed the role of mROS in IL-12 plus IL-18-induced IFN-γ production in naïve and in *in vitro*-differentiated memory-like CD4^+^ T cells, and found significantly higher amounts of mROS and IFN-γ in the later. We demonstrated that mROS signal controls STAT4 and NF-κB activation pathways. Finally, we showed that Fas-deficient CD4^+^ T cells produce more mROS compared to WT in response to IL-12 plus IL-18. This was translated to greatly increased IFN-γ production and to higher proportions of CD44^hi^CD62L^lo^ *lpr* CD4^+^ T cells compared to WT. Our findings establish mROS as key component of TCR-independent IFN-γ production and uncover an unknown role of Fas in regulating mROS and IFN-γ production, providing rationale for targeting mROS in autoimmunity-related hyperinflammation.

## Materials and Methods

### Animals

8-12 weeks old mice were used for CD4^+^ T cell isolation. C57BL/6 mice (WT) were from Harlan Interfauna Ibérica and Fas-deficient C57BL/6-*lpr* mice (*lpr*) were from Jackson laboratories. Mice were kept in SPF conditions. All animal experiments were designed in compliance with European Union and national regulations, and were approved by the CNB Bioethics Committee.

### *In vitro* CD4^+^ T cell activation and memory-like differentiation

Naïve CD4^+^ T cells were purified from mouse spleens using Negative Isolation Kit (Dynal Biotech). Cell purity was >95% as measured by flow cytometry. For cell culture, we used complete media containing 20 ng/ml human recombinant interleukin-2 (rIL-2, PeproTech). Naïve CD4^+^ T cells (10^6^/ml) were stimulated with concavalin A (ConA; 1.5 μg/ml, Sigma) or with IL-12 (10 ng/ml) plus IL-18 (10 ng/ml) for indicated time points. For memory-like differentiation, naïve CD4^+^ T cells were first stimulated with ConA (1.5 μg/ml) for 24h, washed and cultured (0.5 × 10^6^/ml) in presence of 20 ng/ml rIL-2 for 6 days. Differentiated memory-like cells were stimulated with ConA or with IL-12 (10 ng/ml) plus IL-18 (10 ng/ml) for indicated time points. DPI (5 μM; Sigma), rotenone (2.5 μM; Sigma) or antimycin A (4 μM) were added to cell culture 30 min before the stimulus.

### Flow cytometry

For detection of mitochondrial superoxide, cells were incubated with MitoSOX Red (Invitrogen) at 5 μM final concentration for 30 min at 37 °C and stimulated for indicated time points. Cells were then washed with PBS, centrifuged at 300 × *g* for 5 min, and analyzed immediately using flow cytometry. MitoSOX Red fluorescence was detected at the excitation and emission wavelengths of 488 and 585 nm, respectively. To control for baseline fluorescence, samples from each experiment were stained according to the above procedure and left unstimulated. MFI fold induction was calculated by dividing stimulated by unstimulated values.

For measurement of mitochondrial membrane potential TMRM (Tetramethylrhodamine, Methyl Ester, Perchlorate; ThermoFisher) was added to the cells at 30 nM final concentration. After 30 min, cells were analyzed by flow cytometry. Control cells were pre-incubated with 5 μM FCCP for 10 min at 37 °C to depolarize the membrane and show background staining.

Intracellular cytokine staining and cell cycle analysis were performed according to standard procedure described in Supplementary Material. Stained cells were analyzed on Gallios flow cytometer from Beckman Coulter. The data were analyzed using FlowJo software (Tree Star).

### Immunoblotting

Cells were treated as stated and the immunoblotting was performed as described in Supplementary Material.

### EMSA

EMSA was performed as described previously (19) and described in Supplementary Material.

### Statistical Analysis

Statistical significance was determined by unpaired 2-tailed Student’s *t* test for comparisons between two groups, or by 1- or 2-way ANOVA for multiple comparisons, followed by Sidak’s or Tukey post-hoc test. Differences were considered significant when *p*<0.05. All statistical analyses were conducted using Prism 8 software (GraphPad).

## Results

### Memory-like CD4^+^ T cells produce increased mROS and IFN-γ in response to IL-12/IL-18 compared with naïve cells

Naïve CD4^+^ T cells purified from mouse spleens were stimulated *ex vivo* with IL-12/IL-18 to induce IFN-γ production independently of TCR, as previously described (20). Alternatively, T cells were exposed to concavalin A (ConA), which is known for mimicking the physiological TCR cross-linking leading to T cell activation and ROS production (21). After primary TCR activation and IL-2 dependent culture, memory-like CD4^+^ T cells were re-stimulated with either ConA or with IL-12/IL-18. As previously described (22), naïve CD4^+^ T cells, characterized by CD62L^hi^CD44^lo^ phenotype, efficiently differentiated into central and effector memory cell subsets after IL-2 expansion (Supplementary Figure 1). Upon 24 h of TCR stimulation, intracellular staining showed significant but relatively low IFN-γ production both after primary and secondary activation of naïve and memory-like cells, respectively (Figure 1A). IL-12/IL-18 stimulation induced similar IFN-γ levels as ConA in naïve CD4^+^ cells (Figure 1A). By contrast, the proportions of IFN-γ-producing cells after IL-12/IL-18 stimulation were strikingly higher in memory-like compared with naïve cells (Figure 1A). The data showed that IL-12/IL-18 induce IFN-γ more efficiently in memory-like CD4^+^ T cells compared to TCR-dependent stimulation.

**Figure 1.**
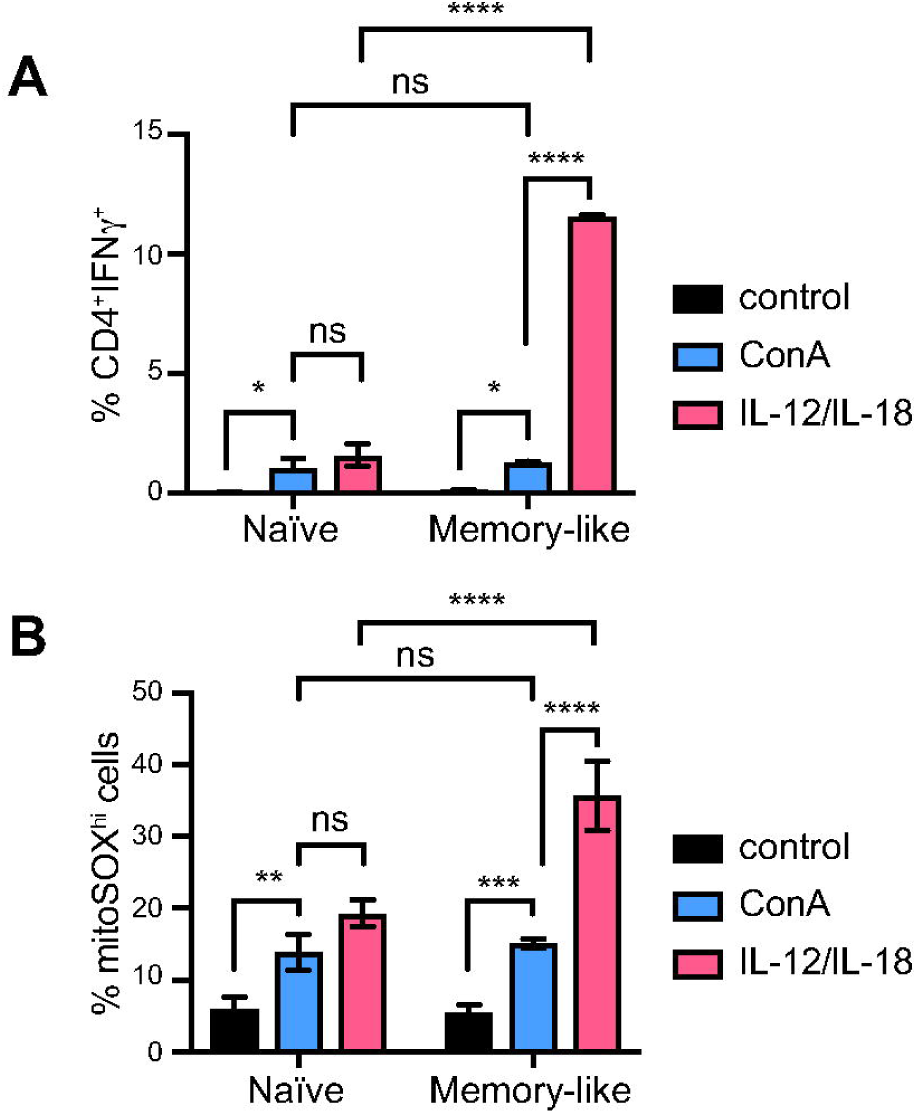
IFN-γ production and mROS levels in memory vs. naïve CD4^+^ T cells. Naïve CD4^+^ T cells were isolated from WT mouse spleens and stimulated with ConA or IL-12 and IL-18. To generate memory T cells, ConA-stimulated CD4^+^ cells were expanded in IL-2 for 6 days and re-stimulated with either ConA or IL-12 and IL-18. **(A)** Flow cytometry analysis of the intracellular staining showing the percentage of IFN-γ-producing CD4^+^ naïve and memory T cells after 24 h stimulation with ConA or IL-12 and IL-18. **(B)** Flow cytometry analysis of mitoSOX red fluorescence showing the percentage of mitoSOX^hi^ cells in naïve and memory cells at 1 hour after ConA or Il-12 and IL-18 stimulation. The graphs show mean ± SD (*n* = 3), \**p*<0.05, \*\**p*<0.01, \*\*\**p*<0.001, \*\*\*\**p*<0.0001, 2-way ANOVA (with Sidak’s correction for multiple comparison).

We measured mitochondrial superoxide production with MitoSOX Red in naïve and memory-like T cells after TCR-dependent and −independent stimulation. After ConA treatment, the percentage of MitoSOX^hi^ cells was significantly increased compared to unstimulated cells, but we found no significant differences in mROS production between naïve and memory-like cells (Figure 1B). In naïve cells, the proportions of MitoSox^hi^ cells after IL-12/IL-18 stimulation were similar to those induced by ConA (Figure 1B). However, IL-12/IL-18 induced significantly higher proportions of MitoSOX^hi^ cells in the memory-like cell population compared to that of naïve cells (Figure 1B). We also measured MitoSOX MFI and obtained similar results (Supplementary Figure 2).

These data showed that memory-like cells displayed enhanced responses after IL-12/IL-18 compared to ConA stimulation in terms of both IFN-γ and mROS production, while the two types of stimulation had similar effects on naïve T cells.

### IL-12/IL-18-dependent IFN-γ production by naïve CD4^+^ T cells requires mROS activation

To investigate whether mROS are functionally related to IL-12/IL-18-induced IFN-γ production by naïve CD4^+^ T cells, we inhibited mROS production using DPI (diphenyleneiodonium) (23, 24). Alternatively, we used specific mitochondrial complex I inhibitor rotenone.

Upon T cell stimulation we detected two cell populations: MitoSOX^lo^ and MitoSOX^hi^ cells. While in MitoSOX^hi^ cell population MitoSOX MFI significantly increased at 30- and 60-min post IL-12/IL-18 or ConA stimulation, MitoSOX^lo^ cells remained unresponsive (Supplementary Figure 3 and data not shown). We therefore focused our analyses on the MitoSOX^hi^ population, which represented activated T cells.

We analyzed early MitoSOX levels with or without mROS inhibitors after T cell activation over a period of two hours. MitoSOX was significantly increased at 30 min post stimulation and remained elevated for at least two hours, as indicated by MitoSOX median fluorescence intensity (MFI) (Figure 2A). At all time points up to 2 h post-stimulation, the activation-induced mROS increase was dramatically suppressed upon DPI treatment, to mROS levels of unstimulated cells (Figure 2A). These data show that DPI inhibits mROS production in cells that are responsive to IL-12/IL-18. Similar results were obtained using rotenone (not shown).

**Figure 2.**
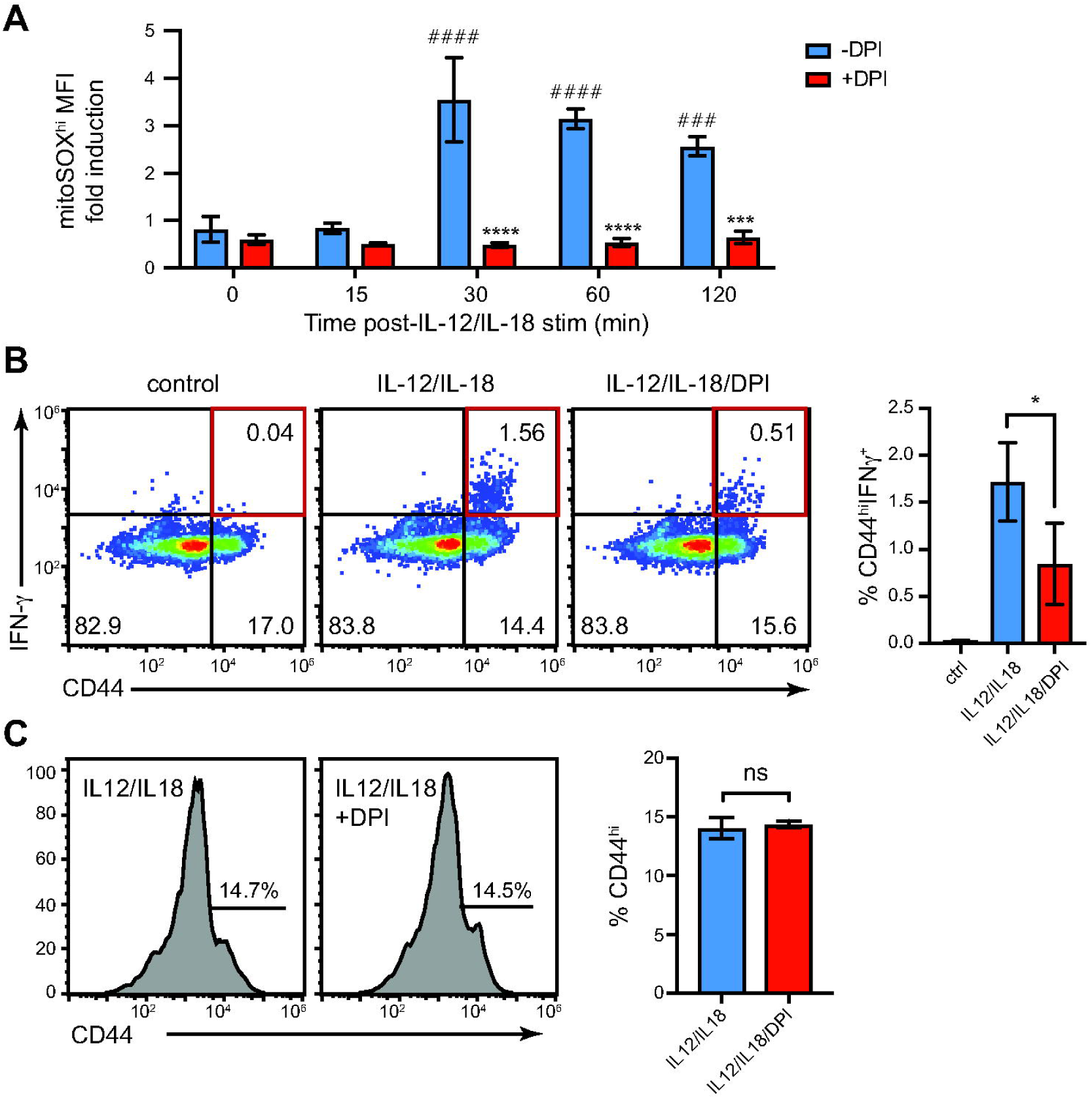
DPI treatment lowers mROS levels and inhibits IFN-γ production induced by IL-12 plus IL-18 in naïve CD4^+^ T cells. Naïve CD4^+^ T cells from WT mouse spleens were stimulated with IL-12 and IL-18 and analyzed by flow cytometry. **(A)** MitoSOX red fluorescence staining showing mROS induction at early time points after IL-12 plus IL-18 stimulation, and its reduction by DPI treatment. Gated on mitoSOX^hi^ population. **(B)** Intracellular staining showing the frequency of IFN-γ-producing CD44^hi^ cells after 24 h of Il-12 plus IL-18 stimulation, and its reduction by DPI treatment. Gated on CD4^+^ cells. **(C)** Similar surface expression of CD44 activation marker in control and DPI-treated cells after 24 h of IL-12 plus IL-18 stimulation. Histograms and dot-plots are representative of at least 3 experiments performed. Graphs show mean ± SD (*n*=3); \**p*<0.05; \*\*\**p*<0.001; \*\*\*\**p*<0.0001; two-way ANOVA with Sidak’s correction; # compared to 0 time point, * DPI treatment compared to control (A); one-way ANOVA with post-hoc Tukey test (B) and (C). *n* represents values obtained from different mice.

IL-12/IL-18 stimulation of naïve T cells induced IFN-γ production in a low cell proportion (∼1.5%) that were also CD44^hi^, at 24h post-stimulation (Figure 2B, left). Importantly, DPI treatment reduced IFN-γ production by these cells (Figure 2B, left) in a statistically significant manner (Figure 2B, right). However, DPI had no effect on the expression of CD44 (Figure 2C). Overall, our results suggest that mROS control IFN-γ production after activation of naïve CD4+ T cells.

### mROS inhibition lowers IFN-γ production by memory-like CD4^+^ T cells and compromises their CD44^hi^ phenotype

Memory-like CD4^+^ T cells generated after primary TCR activation and IL-2-dependent culture significantly increased mROS production as early as 15 min post IL-12/IL-18 stimulation (Figure 3A). The impact of mROS on cytokine-induced signaling is likely more relevant in memory-like than in naïve CD4^+^ T cells, as in the later mROS increase was not detected until 30 min post-stimulation (Figure 2A).

**Figure 3.**
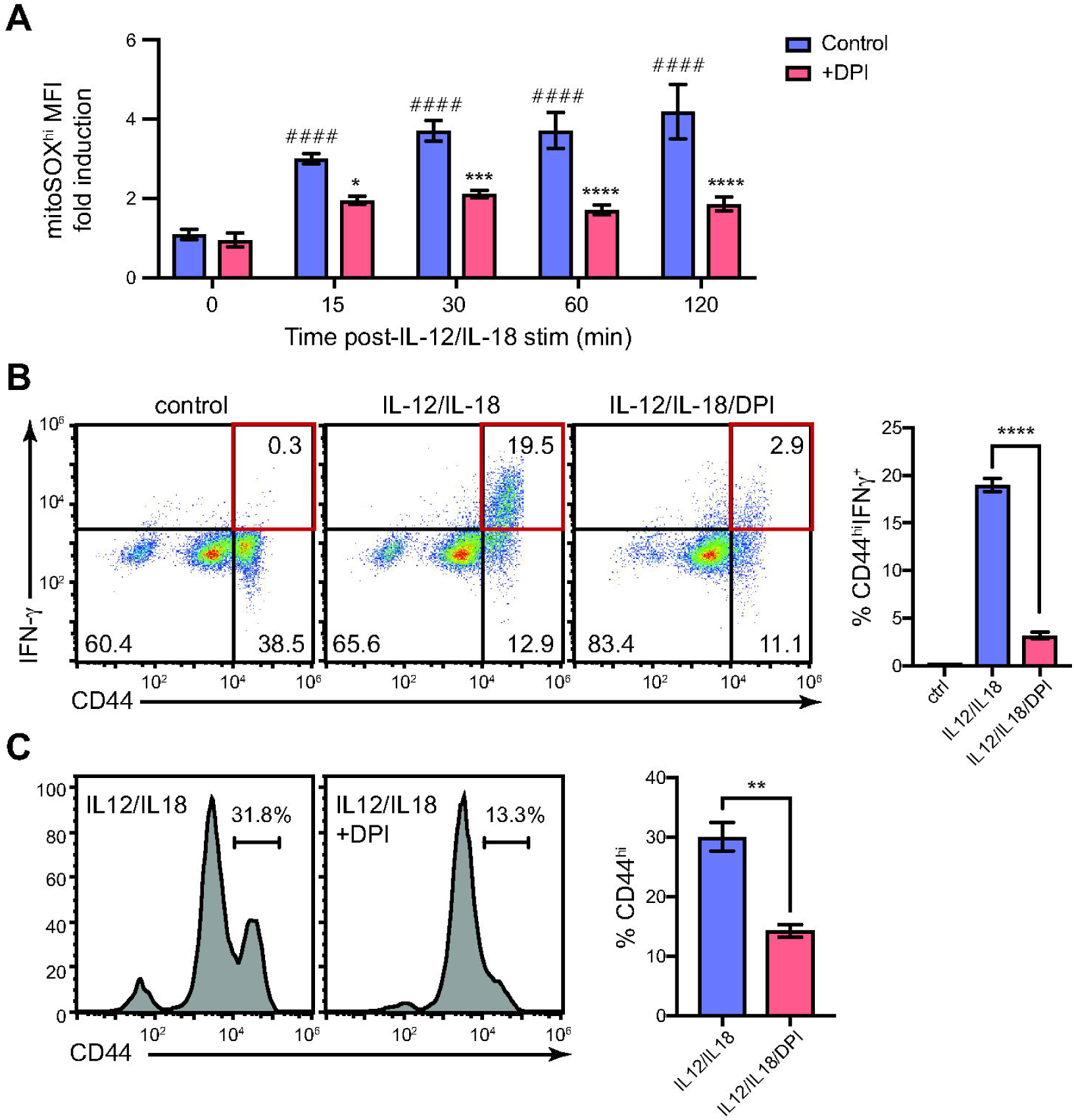
DPI treatment lowers mROS levels and inhibits IFN-γ production in differentiated memory-like CD4^+^ T cells. CD4^+^ T cells from WT mouse spleens were stimulated with ConA for 24 h, expanded in the presence of IL2 for 6 days, and stimulated with IL-12 and IL-18. **(A)** mitoSOX red fluorescence showing increased mROS production in the early time points of IL-12/IL-18 stimulation, and its reduction after DPI treatment. **(B)** Intracellular staining showing reduced frequency of IFN-γ-producing CD44^hi^ effector/memory T cells after DPI treatment; gated on CD4^+^ cells. **(C)** Flow cytometry analysis showing decreased surface expression of CD44 after DPI treatment. Dot plots and histograms are representative of at least 3 experiments performed. Graphs show mean ± SD (*n*=3); \**p*<0.05; \*\**p*<0.01; \*\*\**p*<0.001; \*\*\*\**p*<0.0001, two-way ANOVA with Sidak’s correction; # compared to 0 time point, * DPI treatment compared to control (A); one-way ANOVA with post-hoc Tukey test (B) and (C). *n* represents values obtained from different mice.

DPI led to a prominent reduction of mROS, as detected by MitoSOX at all time points tested after IL-12/IL-18 treatment (Figure 3A). Memory-like cells were more responsive to IL-12/IL-18 stimulation compared to naïve cells, as ∼10% of the total CD4^+^ population were CD44^hi^ cells and positive for IFN-γ at 24 h post-stimulation (Figure 3B). The mROS inhibitor DPI nearly abrogated IFN-γ production in memory-like T cells (Figure 3B, left) in statistically significant manner (Figure 3B, right). In addition, DPI led to significant downregulation of CD44 expression, suggesting that mROS regulates the activation potential of memory-like T cells (Figure 3C). Of note, we didn’t observe any marked effect of DPI treatment on cell viability nor on cell cycle distribution (Supplementary Figure 4A). Our results support the notion that mROS are integral to the maintenance of activated CD44^hi^ phenotype and control the TCR-independent IFN-γ production.

### Validation of DPI as an attenuator of IFN-γ production through inhibition of mROS production

Our results showed that DPI strongly inhibits mROS-dependent IFN-γ production by memory-like CD4^+^ T cells. DPI is also known for inhibiting Nox enzymes (23, 25–27), raising questions of whether the observed effects of DPI on IFN-γ production might be due to mitochondria-unrelated processes. To address this point and validate the effect of mROS generation on IFN-γ production we tested other respiratory chain blockers. Specific complex I inhibitor rotenone significantly decreased IL-12/IL-18-induced mROS production (Figure 4A), as well as the percentages of IFN-γ-producing and CD44^hi^ cells (Figure 4B), thus providing similar results as DPI. We also used antimycin A, an electron transport chain inhibitor acting at complex III. Although antimycin A has been shown to increase mROS production (28), we found that in the present setting it does not affect mROS at early time points post-IL-12/IL-18 stimulation (Figure 4A). In addition, antimycin A was not effective in blocking IFN-γ production nor in reducing CD44 expression, suggesting that an intact electron transport chain is not required for IL-12/IL-18-induced IFN-γ production.

**Figure 4.**
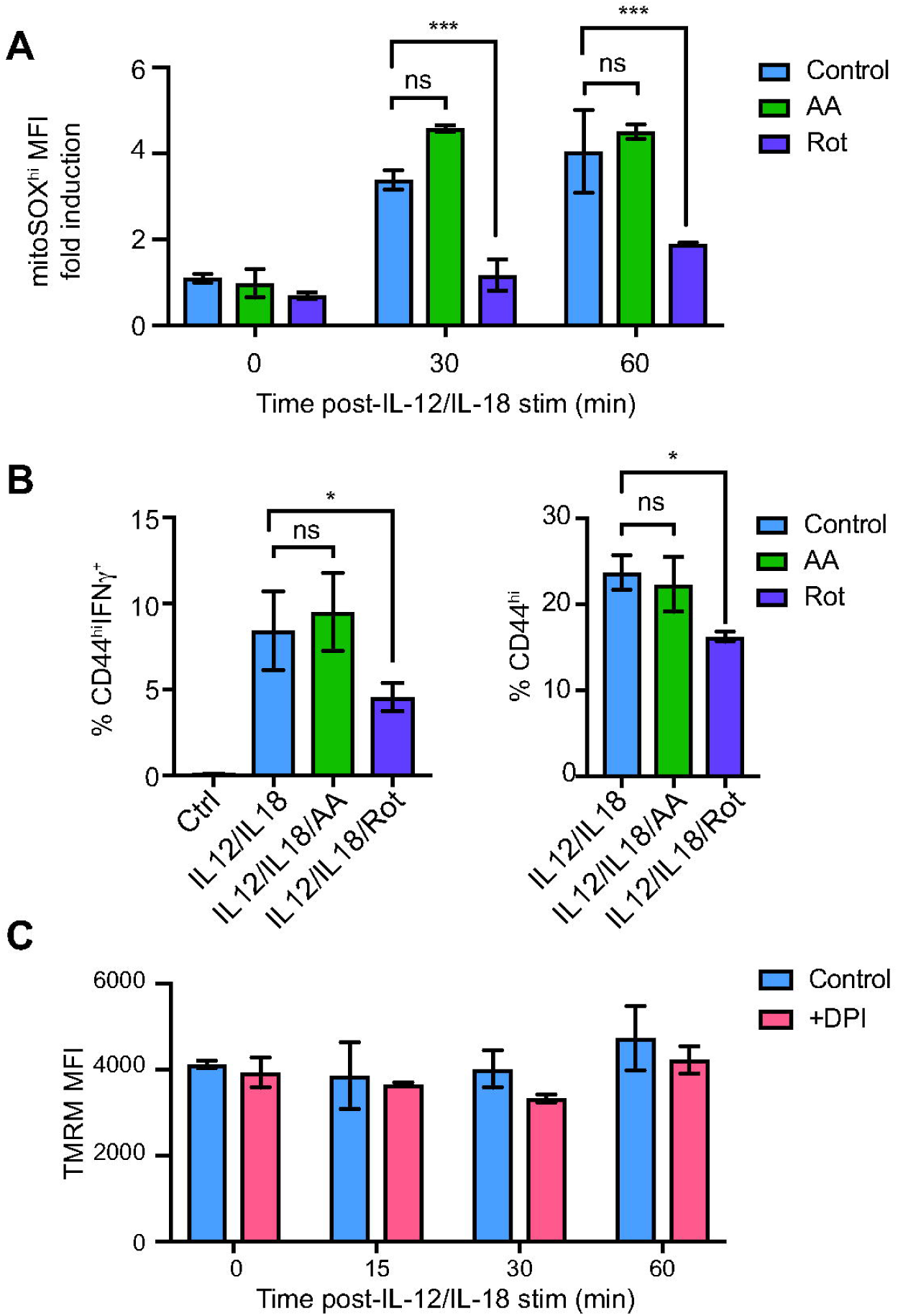
The effect of different respiratory chain blockers on mROS generation, IFN-γ production and mitochondrial membrane potential. CD4^+^ T cells from WT mouse spleens were stimulated with ConA for 24 h, expanded in the presence of IL2 for 6 days, and stimulated with IL-12 and IL-18. **(A)** mROS production in mitoSOX^hi^ population was unaffected by antimycin A and reduced by rotenone treatment. **(B)** Flow cytometry analysis showing the percentages of IFN-γ-producing and CD44^hi^ cells after antimycin A and rotenone treatment; gated on CD4^+^ cells. **(C)** DPI treatment did not affect mitochondrial membrane potential at early time points post IL-12 plus IL-18 treatment, as shown by TMRM fluorescence measured by flow cytometry. FCCP was used to depolarize the membrane and show background staining (not shown). Graphs show mean ± SD (*n*=3); \**p*<0.05; \*\**p*<0.01; \*\*\**p*<0.001; \*\*\*\**p*<0.0001, two-way ANOVA with Sidak’s correction (A) and (C); one-way ANOVA with post-hoc Tukey test (B). *n* represents values obtained from different mice.

Other studies have shown that DPI treatment might affect the overall electron transport activity and mitochondrial membrane potential (29). We therefore used potentiometric TMRM dye to test whether DPI treatment affects the mitochondrial membrane potential during memory-like CD4^+^ T cell activation. TMRM fluorescence showed that neither IL-12/IL-18 nor DPI significantly affected mitochondrial membrane potential at early time points after stimulation (Figure 4C). Thus, it seems that mROS inhibition did not affect the overall electron transport rate, in accordance with previous studies (29).

### mROS-mediated signals activate PKC-θ, AKT, STAT4 and NF-κB

IL-12 and IL-18 act in synergy through different signaling pathways to activate IFN-γ production (30). IL-12 receptor stimulation leads to STAT4 phosphorylation through Jak2 and Tyk2, while IL-18 (also known as IFN-γ-inducing factor) activates the NF-κB pathway via IRAK (interleukin 1 receptor-associated kinase) and TRAF6 (TNF receptor-associated factor 6) (20, 31, 32). We examined the components of these two pathways in memory-like CD4^+^ T cells before and after treatment with mROS inhibitors DPI or rotenone. Before IL-12/IL-18 treatment the NF-κB pathway was kept in its latent state, as seen by high protein levels of IκBα (Figure 5 A, Ctrl). Stimulation induced NF-κB activation, as IκBα levels dropped as early as 15 min and 30 min (highest reduction) post-stimulation (Figure 5A and B). After DPI or rotenone treatment this drop in IκBα was halted, suggesting that NF-κB activation is delayed in conditions of decreased mROS. We also examined the activation of PKC-θ, as it has been previously implicated in TCR-independent NF-κB regulation (33). We found increased PKC-θ phosphorylation as early as 15 min post-IL-12/IL-18 treatment (Figure 5A and B). Both inhibitors reduced the phospho-PKC-θ levels, which was evident at 30 min and more so at 1 h post-treatment. These data suggest that early mitochondrial oxidative signal promotes PKC-θ and NF-κB activation after IL-12/IL-18 signaling. In accordance with this, electromobility shift showed elevated NF-κB nuclear activity at 1h post-IL-12/IL-18 treatment, which was markedly decreased by DPI or rotenone (Figure 5C).

**Figure 5.**
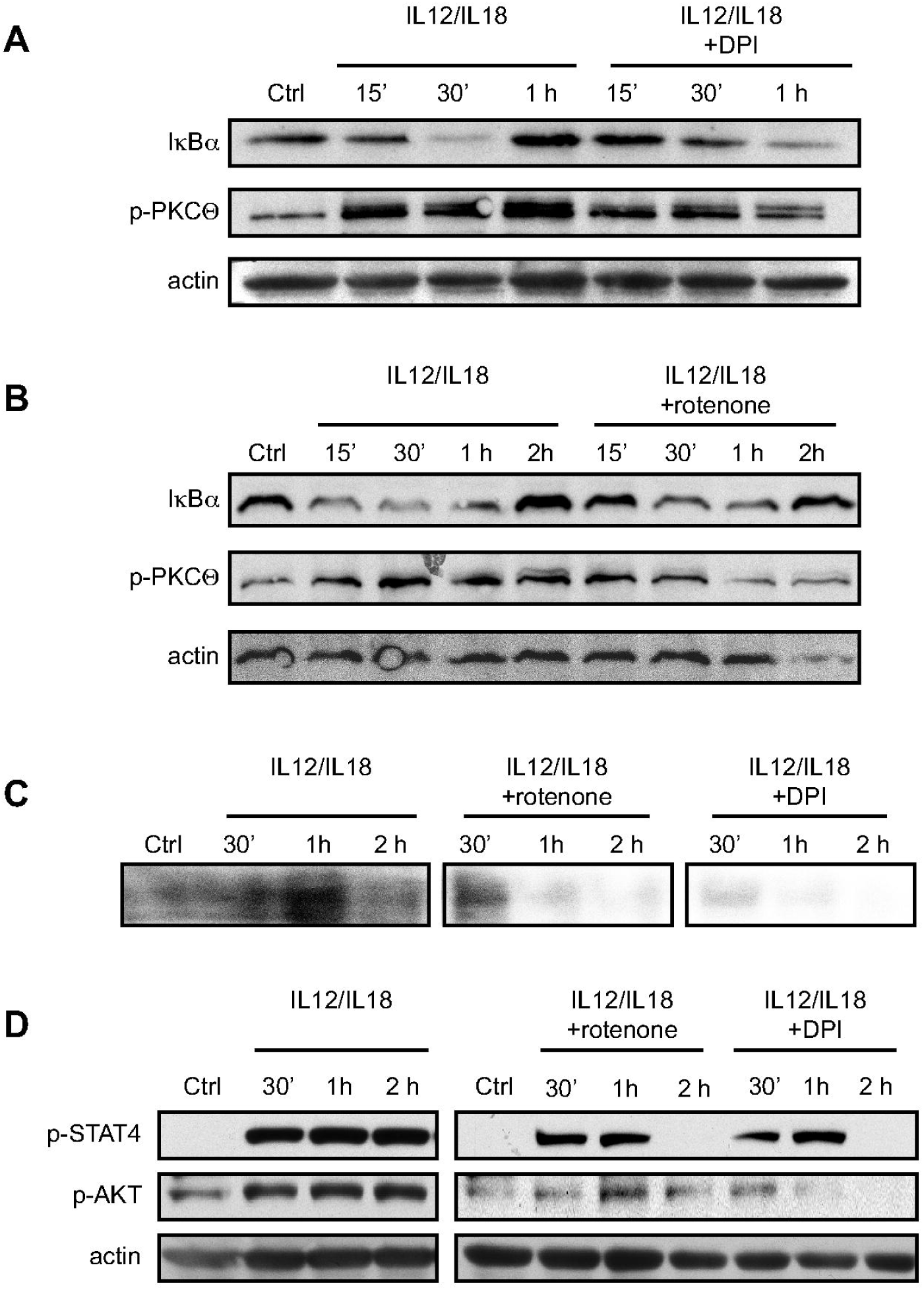
The effect of different inhibitors on signaling pathways triggered by IL-12 and IL-18. CD4^+^ T cells from WT mouse spleens were stimulated with ConA for 24 h, expanded in the presence of IL2 for 6 days, and stimulated with IL-12 and IL-18. Immunoblot analysis showing IκBα and p-PKC-θ and levels after DPI **(A)** and rotenone **(B)** treatment. **(C)** EMSA analysis showing reduced NF-κB nuclear activity after DPI and rotenone treatment. **(D)** Immunoblot analysis of STAT4 and AKT phosphorylation after NAC, rotenone and DPI treatment. β-actin was used as loading control. Shown are representative gels of at least 2 experiments performed.

STAT4 is key transcription factor recruited by IL-12-derived signal to regulate IFN-γ (31). Phosphorylation of STAT4 was evident after IL-12 plus IL-18 stimulation (Figure 5D). Both after rotenone and DPI treatment, STAT4 phosphorylation was markedly decreased early after IL-12/IL-18 stimulation (Figure 5D). This suggested a direct mROS role in IL-12-dependent STAT4 activation. IL-18 in certain cell types activates AKT (34), and mROS affects AKT phosphorylation (35). We detected increased AKT phosphorylation after IL-12/IL-18 treatment (Figure 5D). mROS inhibition using rotenone or DPI markedly decreased phospho-AKT levels (Figure 5D).

Altogether, these data further strengthen the notion that mROS trigger the early signaling events downstream of IL-12 and IL-18 receptors leading to IFN-γ production independently of TCR activation.

### IFN-γ hyperproduction in *lpr* CD44^hi^CD62L^lo^ effector/memory CD4^+^ T cells is related to increased mROS

*lpr* (lymphoproliferation spontaneous mutation) mice are Fas (CD95) deficient and show defective AICD (activation-induced cell death) of *in vitro* re-stimulated T cells (36, 37). *In vivo*, one of the symptoms caused by Fas deficiency in *lpr* mice is the accumulation of effector/memory CD44^hi^CD62L^lo^ CD4^+^ T cells and their intrinsic hyperactivation accompanied by hyperproduction of IFN-γ in response to IL-12/IL-18 (13). To investigate the role of mROS in the responses of Fas-deficient memory-like cells, we purified CD4^+^ T cells from WT and *lpr* mice and, after initial ConA stimulation and IL-2 expansion, we stimulated the cells with IL-12/IL-18. Significantly higher mROS levels were induced in *lpr* compared with WT cells, as seen by MitoSOX at 60 min after stimulation (Figure 6A). The mROS production was greatly diminished by DPI in both WT and *lpr* cells (Figure 6B). Of note, the proportions of CD44^hi^ cells were similar between WT and *lpr*, suggesting that Fas deficiency did not cause increased accumulation of memory-like cells under these experimental conditions and TCR-independent stimulation (Figure 6C). Accordingly, cell cycle analysis showed no apparent difference in apoptosis or proliferation in WT and *lpr* CD4^+^ T cells after IL-12 plus IL-18 stimulation (Supplementary Figure 4B). Instead, *lpr* CD44^hi^ T cells were hyperactivated, as seen by significantly increased IFN-γ production compared to WT cells (∼20 *vs*. 30% of the total CD4^+^ population) at 24 h post-IL-12/IL-18 (Figure 6D). DPI treatment significantly reduced the IFN-γ production in both WT and *lpr* CD44^hi^ T cells (Figure 6D). These results suggest that the increased mROS production drives the increased IFN-γ production by *lpr* CD44^hi^ T cells.

**Figure 6.**
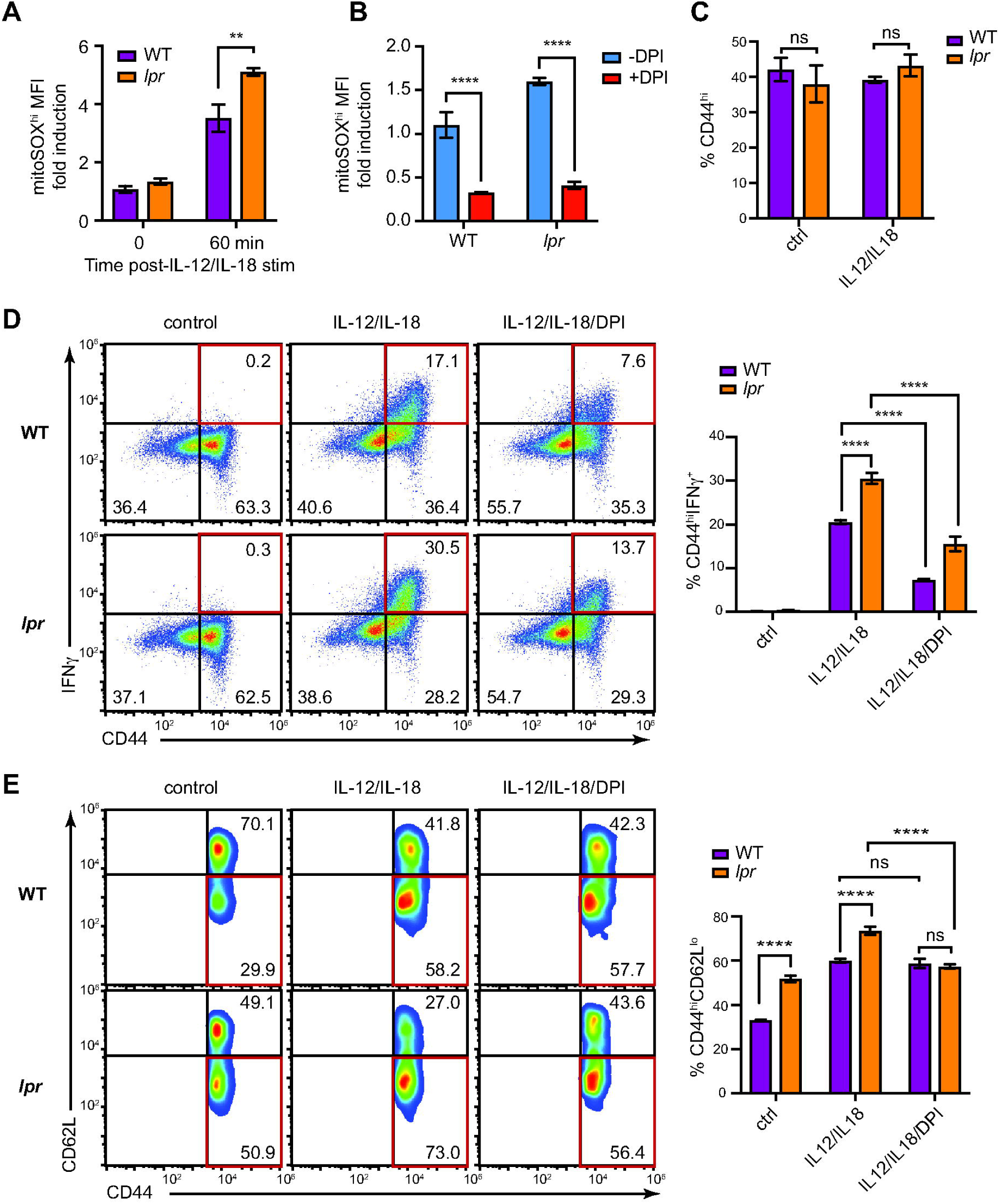
mROS drives increased IFN-γ production in *lpr* CD44^hi^CD62L^lo^CD4^+^ T cells. CD4^+^ T cells from WT and *lpr* mouse spleens were stimulated with ConA for 24 h, expanded in presence of IL-2 for 6 days and stimulated with IL-12 and IL-18. **(A)** Flow cytometry analysis showing increased mROS production in mitoSOX^hi^ *lpr* cells compared with WT at 60 min after IL-12 plus IL-18 stimulation. **(B)** The effect of DPI treatment on mROS production by WT and *lpr* cells at 60 min after IL-12 plus IL-18 stimulation. **(C)** Flow cytometry analysis showing unaffected CD44 surface expression in WT and *lpr* cells after IL-12 plus IL-18 treatment. **(D)** Intracellular staining showing increased frequency of IFN-γ-producing CD44^hi^ *lpr* cells compared with WT and their reduction after DPI treatment. **(E)** Flow cytometry analysis of CD62L and CD44 surface expression showing the increased proportions of *lpr* CD44^hi^C62L^lo^ cells compared with WT and their reduction after DPI treatment. Shown are representative dot plots of 3 experiments performed. Graphs show mean ± SD (*n*=3); \**p*<0.05; \*\**p*<0.01; \*\*\**p*<0.001; \*\*\*\**p*<0.0001, two-way ANOVA with Sidak’s correction. *n* represents values obtained from different mice.

*lpr* CD4^+^ T cells showed increased proportions of CD44^hi^/CD62L^lo^ effector/memory cells compared with the WT (∼30 *vs*. 50%), which were further increased after IL-12/IL-18 treatment (∼58 *vs*. 73%, Figure 6E). Although DPI did not affect CD62L^hi^ and CD62L^lo^ proportions in WT cells, it significantly reduced the population of *lpr* CD44^hi^/CD62L^lo^ effector/memory cells, to the levels comparable to those in the WT cell population (Figure 6E). Altogether, our results corroborate that the increased mROS production drives not only the hyperactivation and increased potential for IFN-γ production in Fas-deficient CD4^+^ T cells, but also their intensified differentiation towards the effector/memory CD44^hi^/CD62L^lo^ phenotype. This points to Fas as a regulator of mitochondrial ROS that ultimately controls the status of memory-like T cells independently of the proapoptotic role of Fas in such cells.

## Discussion

It is established that TCR engagement induces mitochondrial activation and mROS production that modulates the redox state of signaling molecules governing the early events downstream of TCR induction. Here, we analyzed the role of mROS in recall responses of memory-like T cells. mROS induction was similar in naïve and memory-like cells after TCR-dependent signaling, while in response to IL-12 plus IL-18 memory-like CD4^+^ T cells showed a great and highly significant increase in mROS and IFN-γ production compared to naïve T cells. This suggested that memory-like CD4^+^ T cells stimulated by IL-12 plus IL-18 acquired a singular capacity in producing mROS, and our study was geared at examining the effect of such TCR-independent triggering. We report the following major findings: 1) Inhibition of the increased mROS production by memory-like T cells after exposure to IL-12 plus IL-18 was directly associated to IFN-γ production and to maintaining the memory-like phenotype of stimulated cells; 2) The mechanism that drives IFN-γ production is regulated by the effect of mROS on the activation of the pathways that depend on IL-12 plus IL-18 signaling; 3) Our results uncovered a previously unknown Fas role as a regulator of mROS induction, since *lpr* memory-like CD4^+^ T cells produced higher mROS levels compared to WT cells, leading to elevated IFN-γ and increased differentiation to effector/ memory T cells.

TCR stimulation of CD4^+^ cells triggers the early production of mROS, which act as secondary messengers in regulating IL-2 production and cell proliferation (6, 38). Additionally, at later stages of CD4^+^ T cells stimulation, mitochondrial activity and mROS production are increased (39). Mitochondria activation is associated with enhanced recall responses of memory CD8^+^ T cells after TCR-dependent restimulation, but a possible mROS association with memory T cell responses was not assessed (40). Nevertheless, cytosolic ROS was found increased in *in vitro* memory-like CD4^+^ T cells after TCR dependent stimulation (5). In our study we found similar mROS levels in naïve and memory-like CD4^+^ T cells early after TCR activation. We used IL-12 plus IL-18 stimulation to investigate the role of mROS in memory-like CD4^+^ T cells, a protocol that did not induce T cell apoptosis or proliferation, but greatly increased mROS in memory-like compared to naïve T cells after activation. Overall, our findings suggest that mROS production in T cells depends on their activation state and the stimulus type.

mROS levels were directly linked to the hyperactivation status of T cells in response to IL-12 plus IL-18 induction, as inhibition of mROS production by DPI or Rotenone reflected a greatly lowered IFN-γ expression and the diminished expression of the CD44 activation/memory marker. Apart from mROS, other aspects of mitochondrial activity are needed for correct T cell function and TCR-induced activation. As part of the immunological synapse, mitochondria participate in Ca^2+^ uptake and thus determine amplitude and duration of Ca^2+^ signal. Mitochondrial uncoupling results in inhibition of the TCR-induced Ca^2+^ signal due to diminished mitochondrial membrane potential and reduced ability to prolong Ca influx (41). In our approach the mitochondrial membrane potential remained unaffected by mROS inhibition, indicating that mROS is the promotor of memory-T cell hyperactivation.

IFN-γ production is induced through synergy of signaling driven by IL-12R and IL-18R stimulation pathways (30). mROS inhibition reduced IL-12R-dependent STAT4 phosphorylation and IL-18-dependent NF-κB activation, indicating that mROS drive signaling of both pathways. Kinetic analysis showed that mROS affects these pathways early on after exposure of cells to IL-12 plus IL-18. Other molecules such as PKC-θ or AKT linked to these pathways showed increased phosphorylation due to increased mROS. The redox conditions that are associated to increased mROS production, could be responsible for alteration in phosphorylation patterns, as kinase and phosphatase activities are redox-dependent. Whether the increased mROS activate specific pathway components or have a multiple effect in the phosphorylation and activation remains to be explored. Another important issue is to understand the reasons behind the specificity of mROS production after secondary IL-12 plus IL-18 stimulation. We hypothesize that components of the IL-12R or the IL-18R signaling pathways may drive ETC respiration. Indeed, PKC-θ (28) is shown to interact with and activate mitochondria, and Tyk2 that forms part of the Jak family and a component of the IL-12R signaling regulates mitochondrial activation (42).

Compared to normal T cells, *lpr* memory-like CD4^+^ overproduced mROS and IFN-γ after IL-12 plus IL-18 activation. Inhibition of mROS induction reduced not only the elevated IFN-γ but also the CD44^hi^/CD62L^lo^ effector/memory cells that are responsible for the *in vivo* IFN-γ and lupus-like symptoms of *lpr* mice (13). Thus, our data suggest that Fas through non-apoptotic signaling (IL-12/IL-18) limits the production of mROS,, compromising differentiation into effector/memory T cells and IFN-γ overproduction. This might provide a feedback mechanism to control T cell homeostasis under chronic inflammation and prevent development of autoimmunity. Therefore, uncontrolled mROS production by T cells in *lpr* mice or humans with ALPS (autoimmune lympho-proliferative syndrome) (43) might favor lymphadenopathy development, as mROS inhibition lowers Fas-dependent AICD (28). Other non-apoptotic Fas roles have been described, especially after primary T cell stimulation (44). Further studies are clearly needed for the better understanding of the mechanism that would explain the effect of Fas on mROS regulation. Overall, the results unveil a previously unknown apoptosis-independent role of Fas in regulating mROS production and associated hyperinflammatory outcomes. Our findings identify mROS as a driver of inflammation in *lpr* disease and potentially other autoimmune disorders. These observations strongly suggest that blockade of mROS provides a therapeutic strategy to decrease inflammation in T cell-mediated inflammatory disorders.

## Acknowledgements

This work was funded by the grant from Comunidad de Madrid (CAM; B2017/BMD-3804) to D. Balomenos. G. Rackov holds postdoctoral Juan de la Cierva Fellowship. None of the funding institutions had the role in the design of the study and collection, analysis and interpretation of the data and writing of the manuscript.

## Conflict of interest

The authors declare no competing financial interest in relation to the work described.

## Availability of data and materials

All data generated and analyzed during this study are included in this article.

## Authors’ contributions

DB and GR conceived and designed the study, analyzed the data and wrote the manuscript. CMA and MAM provided input in designing the project. GR, PTZ, SCP and RS performed the experiments. All authors read, discussed and approved the manuscript.

## Abbreviations used in this article

AICD: activation-induced cell death
ALPS: autoimmune lympho-proliferative syndrome
ConA: concavalin A
DPI: diphenyleneiodonium
ETC: electron transport chain
FCCP: carbonyl cyanide 4-(trifluoromethoxy)phenylhydrazone
IRAK: interleukin 1 receptor-associated kinase
MFI: median fluorescence intensity
mROS: mitochondrial reactive oxygen species
Nox: NADPH oxidase
TMRM: tetramethylrhodamine, methyl ester, perchlorate
TRAF6: TNF receptor-associated factor 6
WT: wild-type

## Notes

### Competing Interest Statement

The authors have declared no competing interest.

## References

1. Tarasov, A. I., E. J. Griffiths, and G. A. Rutter. 2012. Regulation of ATP production by mitochondrial Ca(2+). Cell Calcium 52: 28–35.

2. Koopman, W. J. H., L. G. J. Nijtmans, C. E. J. Dieteren, P. Roestenberg, F. Valsecchi, J. A. M. Smeitink, and P. H. G. M. Willems. 2010. Mammalian mitochondrial complex I: biogenesis, regulation, and reactive oxygen species generation. Antioxid. Redox Signal. 12: 1431–70.

3. Desdín-Micó, G., G. Soto-Heredero, and M. Mittelbrunn. 2018. Mitochondrial activity in T cells. Mitochondrion 41: 51–57.

4. Han, D., F. Antunes, R. Canali, D. Rettori, and E. Cadenas. 2003. Voltage-dependent anion channels control the release of the superoxide anion from mitochondria to cytosol. J. Biol. Chem. 278: 5557–63.

5. Kaminski, M. M., S. W. Sauer, C.-D. Klemke, D. Süss, J. G. Okun, P. H. Krammer, and K. Gülow. 2010. Mitochondrial reactive oxygen species control T cell activation by regulating IL-2 and IL-4 expression: mechanism of ciprofloxacin-mediated immunosuppression. J. Immunol. 184: 4827–4841.

6. Sena, L. A., S. Li, A. Jairaman, M. Prakriya, T. Ezponda, D. A. Hildeman, C. R. Wang, P. T. Schumacker, J. D. Licht, H. Perlman, P. J. Bryce, and N. S. Chandel. 2013. Mitochondria Are Required for Antigen-Specific T Cell Activation through Reactive Oxygen Species Signaling. Immunity 38: 225–236.

7. Trinchieri, G. 1994. Interleukin-12: a cytokine produced by antigen-presenting cells with immunoregulatory functions in the generation of T-helper cells type 1 and cytotoxic lymphocytes. Blood 84: 4008–27.

8. Okamura, H., S. Kashiwamura, H. Tsutsui, T. Yoshimoto, and K. Nakanishi. 1998. Regulation of interferon-y production by IL-12 and IL-18. Curr Opin Immunol 10: 259– 264.

9. Yang, J., H. Zhu, T. L. Murphy, W. Ouyang, and K. M. Murphy. 2001. IL-18-stimulated GADD45β required in cytokine-induced, but not TCR-induced, IFN-γ production. Nat. Immunol. 2: 157–164.

10. Munk, R. B., K. Sugiyama, P. Ghosh, C. Y. Sasaki, L. Rezanka, K. Banerjee, H. Takahashi, R. Sen, and D. L. Longo. 2011. Antigen-independent IFN-γ production by human naïve CD4+ T cells activated by IL-12 plus IL-18. PLoS One 6: 1–8.

11. Yu, J. J., C. S. Tripp, and J. H. Russell. 2003. Regulation and Phenotype of an Innate Th1 Cell: Role of Cytokines and the p38 Kinase Pathway. J. Immunol. 171: 6112–6118.

12. Sattler, A., U. Wagner, M. Rossol, J. Sieper, P. Wu, A. Krause, W. A. Schmidt, S. Radmer, S. Kohler, C. Romagnani, and A. Thiel. 2009. Cytokine-induced human IFN-γ-secreting effector-memory Th cells in chronic autoimmune inflammation. Blood 113: 1948–1956.

13. Daszkiewicz, L., C. Vázquez-Mateo, G. Rackov, A. Ballesteros-Tato, K. Weber, A. Madrigal-Avilés, M. Di Pilato, A. Fotedar, R. Fotedar, J. M. Flores, M. Esteban, C. Martínez-A, and D. Balomenos. 2015. Distinct p21 requirements for regulating normal and self-reactive T cells through IFN-γ production. Sci. Rep. 5: 7691.

14. Papadakis, K. A., D. Zhu, J. L. Prehn, C. Landers, A. Avanesyan, G. Lafkas, and S. R. Targan. 2005. Dominant Role for TL1A/DR3 Pathway in IL-12 plus IL-18-Induced IFN-γ Production by Peripheral Blood and Mucosal CCR9 + T Lymphocytes. J. Immunol. 174: 4985–4990.

15. Balasubramani, A., Y. Shibata, G. E. Crawford, A. S. Baldwin, R. D. Hatton, and C. T. Weaver. 2010. Modular utilization of distal cis-regulatory elements controls Ifng gene expression in T cells activated by distinct stimuli. Immunity 33: 35–47.

16. Yang, J., T. L. Murphy, W. Ouyang, and K. M. Murphy. 1999. Induction of interferon-γ production in Th1 CD4+ T cells: Evidence for two distinct pathways for promoter activation. Eur. J. Immunol. 29: 548–555.

17. Gergely, P., B. Niland, N. Gonchoroff, R. Pullmann, P. E. Phillips, and A. Perl.2002. Persistent mitochondrial hyperpolarization, increased reactive oxygen intermediate production, and cytoplasmic alkalinization characterize altered IL-10 signaling in patients with systemic lupus erythematosus. J. Immunol. 169: 1092–101.

18. Perl, A. 2013. Oxidative stress in the pathology and treatment of systemic lupus erythematosus. Nat. Rev. Rheumatol. 9: 674–686.

19. Rackov, G., E. Hernández-Jiménez, R. Shokri, L. Carmona-Rodríguez, S. Mañes, M. Álvarez-Mon, E. López-Collazo, C. Martínez-A, and D. Balomenos. 2016. p21 mediates macrophage reprogramming through regulation of p50-p50 NF-κB and IFN-β. J. Clin. Invest. 126: 3089–103.

20. Robinson, D., K. Shibuya, A. Mui, F. Zonin, E. Murphy, T. Sana, S. B. Hartley, S. Menon, R. Kastelein, F. Bazan, and A. O. Garra. 1997. IGIF Does Not Drive Th1 Development but Synergizes with IL-12 for Interferon-g Production and Activates IRAK and NFkB. Immunity 7: 571–581.

21. Pani, G., R. Colavitti, S. Borrello, and T. Galeotti. 2000. Endogenous oxygen radicals modulate protein tyrosine phosphorylation and JNK-1 activation in lectin-stimulated thymocytes. Biochem. J. 347: 173–181.

22. Cruz, A. C., M. Ramaswamy, C. Ouyang, C. A. Klebanoff, P. Sengupta, T. N. Yamamoto, F. Meylan, S. K. Thomas, N. Richoz, R. Eil, S. Price, R. Casellas, V. K. Rao, J. Lippincott-Schwartz, N. P. Restifo, and R. M. Siegel. 2016. Fas/CD95 prevents autoimmunity independently of lipid raft localization and efficient apoptosis induction. Nat. Commun. 7.

23. Li, Y., and M. A. Trush. 1998. Diphenyleneiodonium, an NAD(P)H oxidase inhibitor, also potently inhibits mitochondrial reactive oxygen species production. Biochem. Biophys. Res. Commun. 253: 295–299.

24. Bulua, A. C., A. Simon, R. Maddipati, M. Pelletier, H. Park, K.-Y. Kim, M. N. Sack, D. L. Kastner, and R. M. Siegel. 2011. Mitochondrial reactive oxygen species promote production of proinflammatory cytokines and are elevated in TNFR1-associated periodic syndrome (TRAPS). J. Exp. Med. 208: 519–33.

25. Vendrov, A. E., N. R. Madamanchi, Z. S. Hakim, M. Rojas, and M. S. Runge. 2006. Thrombin and NAD(P)H oxidase-mediated regulation of CD44 and BMP4-Id pathway in VSMC, restenosis, and atherosclerosis. Circ. Res. 98: 1254–1263.

26. Cross, A. R., and O. T. Jones. 1986. The effect of the inhibitor diphenylene iodonium on the superoxide-generating system of neutrophils. Specific labelling of a component polypeptide of the oxidase. Biochem. J. 237: 111–116.

27. Devadas, S., L. Zaritskaya, S. G. Rhee, L. Oberley, and M. S. Williams. 2002. Discrete generation of superoxide and hydrogen peroxide by T cell receptor stimulation: selective regulation of mitogen-activated protein kinase activation and fas ligand expression. J. Exp. Med. 195: 59–70.

28. Kaminski, M., M. Kiessling, D. Süss, P. H. Krammer, and K. Gülow. 2007. Novel role for mitochondria: protein kinase Ctheta-dependent oxidative signaling organelles in activation-induced T-cell death. Mol. Cell. Biol. 27: 3625–3639.

29. Lambert, A. J., J. A. Buckingham, H. M. Boysen, and M. D. Brand. 2008. Diphenyleneiodonium acutely inhibits reactive oxygen species production by mitochondrial complex I during reverse, but not forward electron transport. Biochim. Biophys. Acta -Bioenerg. 1777: 397–403.

30. Nakahira, M., H.-J. Ahn, W.-R. Park, P. Gao, M. Tomura, C.-S. Park, T. Hamaoka, T. Ohta, M. Kurimoto, and H. Fujiwara. 2002. Synergy of IL-12 and IL-18 for IFN-gamma gene expression: IL-12-induced STAT4 contributes to IFN-gamma promoter activation by up-regulating the binding activity of IL-18-induced activator protein 1. J. Immunol. 168: 1146–1153.

31. Bacon, C. M., E. F. Petricoin, J. R. Ortaldo, R. C. Rees, A. C. Larner, J. A. Johnston, and J. J. O’Shea. 1995. Interleukin 12 induces tyrosine phosphorylation and activation of STAT4 in human lymphocytes. Proc. Natl. Acad. Sci. U. S. A. 92: 7307– 11.

32. Dinarello, C. A., D. Novick, S. Kim, and G. Kaplanski. 2013. Interleukin-18 and IL-18 binding protein. Front. Immunol. 4: 1–10.

33. So, T., and M. Croft. 2012. Regulation of the PKCΘ-NF-κB axis int lymphocytes by the tumor necrosis factor receptor family member OX40. Front. Immunol. 3: 1–8.

34. Rex, D. A. B., N. Agarwal, T. S. K. Prasad, R. K. Kandasamy, Y. Subbannayya, and S. M. Pinto. 2020. A comprehensive pathway map of IL-18-mediated signalling. J. Cell Commun. Signal. 14: 257–266.

35. Kim, J. H., T. G. Choi, S. Park, H. R. Yun, N. N. Y. Nguyen, Y. H. Jo, M. Jang, J. Kim, J. Kim, I. Kang, J. Ha, M. P. Murphy, D. G. Tang, and S. S. Kim. 2018. Mitochondrial ROS-derived PTEN oxidation activates PI3K pathway for mTOR-induced myogenic autophagy. Cell Death Differ. 25: 1921–1937.

36. Singer, G. G., A. C. Carrera, A. Marshak-Rothstein, C. Martinez, and A. K. Abbas. 1994. Apoptosis, Fas and systemic autoimmunity: the MRL-lpr/lpr model. Curr Opin Immunol 6: 913–920.

37. Walker, L. S., and A. K. Abbas. 2002. The enemy within: keeping self-reactive T cells at bay in the periphery. Nat Rev Immunol 2: 11–19.

38. Murphy, M. P., and R. M. Siegel. 2013. Mitochondrial ROS fire up T cell activation. Immunity 38: 201–2.

39. Akkaya, B., A. S. Roesler, P. Miozzo, B. P. Theall, J. Al Souz, M. G. Smelkinson, J. Kabat, J. Traba, M. N. Sack, J. A. Brzostowski, M. Pena, D. W. Dorward, S. K. Pierce, and M. Akkaya. 2018. Increased Mitochondrial Biogenesis and Reactive Oxygen Species Production Accompany Prolonged CD4 + T Cell Activation. J. Immunol. 201: 3294–3306.

40. Van Der Windt, G. J. W., D. O’Sullivan, B. Everts, S. C. C. Huang, M. D. Buck, J. D. Curtis, C. H. Chang, A. M. Smith, T. Ai, B. Faubert, R. G. Jones, E. J. Pearce, and E. L. Pearce. 2013. CD8 memory T cells have a bioenergetic advantage that underlies their rapid recall ability. Proc. Natl. Acad. Sci. U. S. A. 110: 14336–14341.

41. Hoth, M., D. C. Button, and R. S. Lewis. 2000. Mitochondrial control of calcium-channel gating: A mechanism for sustained signaling and transcriptional activation in T lymphocytes. Proc. Natl. Acad. Sci. 97: 10607–10612.

42. Potla, R., T. Koeck, J. Wegrzyn, S. Cherukuri, K. Shimoda, D. P. Baker, J. Wolfman, S. M. Planchon, C. Esposito, B. Hoit, J. Dulak, A. Wolfman, D. Stuehr, and A. C. Larner. 2006. Tyk2 tyrosine kinase expression is required for the maintenance of mitochondrial respiration in primary pro-B lymphocytes. Mol. Cell. Biol. 26: 8562– 8571.

43. Rieux-Laucat, F., F. Le Deist, C. Hivroz, I. A. Roberts, K. M. Debatin, A. Fischer, and J. P. de Villartay. 1995. Mutations in Fas associated with human lymphoproliferative syndrome and autoimmunity. Science 268: 1347–1349.

44. Balomenos, D., R. Shokri, L. Daszkiewicz, C. Vázquez-Mateo, and C. Martínez-A. 2017. On How Fas Apoptosis-Independent Pathways Drive T Cell Hyperproliferation and Lymphadenopathy in lpr Mice. Front. Immunol. 8.

