## Supplementary Material for "Mitochondrial oxidative stress drives IL-12/IL-18-induced IFN-γ production by CD4^+^ T cells and is controlled by Fas"

**Running title:** mROS and cytokine-induced IFN-γ in memory-like CD4$^+$ T cells

**Key words:** mitochondrial reactive oxygen species, IFN-γ, IL-12/IL-18, Fas, effetor/memory CD4 T cells

***Corresponding authors:**

Dr. Dimitrios Balomenos, Department of Immunology and Oncology, Centro Nacional de Biotecnología-Consejo Superior de Investigaciones Científicas (CNB-CSIC), Calle Darwin 3, Campus de Cantoblanco, 28049 Madrid, Spain. E-mail address:

Dr. Gorjana Rackov, Department of Immunology and Oncology, Centro Nacional de Biotecnología-Consejo Superior de Investigaciones Científicas (CNB-CSIC), Calle Darwin 3, Campus de Cantoblanco, 28049 Madrid, Spain. E-mail address:

### Supplementary Materials and Methods

##### Composition of cell culture complete media

RPMI 1640 containing 50 U/ml penicillin, 100 μg/ml streptomycin, 50 μM 2-mercaptoethanol, 2 mM L-glutamine, 0.1 mM non-essential amino acids, 10 mM HEPES, 1mM sodium pyruvate, 10% Fetal bovine serum.

##### Flow cytometry

To verify purity, isolated cells were stained with anti-CD4-FITC, -CD8-PE and -B220-APC (Beckman Coulter). For naïve and effector/memory phenotyping, $CD4^+$ T cells were stained with anti-CD4-PeCy7, -CD44-APC and -CD62L-PE (Beckman Coulter).

##### Intracellular cytokine staining

To measure intracellular levels of IFN-γ, cells were stimulated for 24 hours and 1 x Brefeldin A (BioLegend) was added to cell culture during the last 3 h of stimulation. Cells were stained with violet LIVE/DEAD stain kit (Invitrogen) for dead cell exclusion. After surface marker staining with anti-CD4-PeCy7 (GK1.5; BioLegend), -CD44-APC, -CD62L-PE (11-7311-82; eBioscience), cells were washed with PBS, fixed and permeabilized with Citofix/Citoperm kit (BD Biosciences) and blocked with 1μl of Fc-receptor-binding antibody (082732121, Beckman Coulter) for 15 min at 4 °C. Cells were stained intracellularly with anti-IFN-γ-FITC (1/100 dilution, eBioscience) and analyzed on a Gallios cytometer (Beckman Coulter).

##### Cell cycle analysis

To analyze cell cycle, $10^6$ cells were permeabilized with detergent, stained with PI according to the manufacturer's instructions (DNA-Prep Reagent Kit, Beckman Coulter) for 30 min at 37 °C, and analyzed by flow cytometry.

##### Immunoblotting

Cells were treated as stated, washed with ice-cold PBS and lysed in a buffer containing 0.2% Nonidet P-40, 10 mM Tris-HCl [pH 7.5], 150 mM NaCl, and 1 x protease and phosphatase inhibitor cocktail (Roche), for 20 min at 4 °C. 20 μg of protein was resolved per lane in 12% SDS-PAGE, transferred to a nitrocellulose membrane (75 min, 300 mA) and immunoblotted using antibodies against phospho-STAT4 (Sc-101804) from Santa Cruz Biotechnology; phospho-AKT (#3787), phospho-PKC-θ (#9377) and IκBα (#9242) from Cell Signaling; and actin (AC-15) from Sigma. HRP-conjugated secondary antibodies (Dako) were used at 1:2,000 dilution for 1h at room temperature. Blots were visualized using Western Lightning Plus-ECL (PerkinElmer).

##### EMSA

Cells were washed with ice-cold PBS and nuclear extracts were obtained using nuclear/cytosol Fractionation kit (BioVision). Double stranded NF-κB consensus oligonucleotides (5'-AGT TGA GGG GAC TTT CCC AGG C-3') were obtained from

Promega and end-labeled with [$\gamma^{32}$P]ATP (PerkinElmer) using T4 polynucleotide kinase (Promega). In a 25 μl reaction volume, 5 μg of nuclear extract was incubated with 0.5 ng labeled oligonucleotide probe, 10 mM HEPES, 1 mM $MgCl_2$, 35 mM NaCl, 0.5 mM EDTA, 0.5 mM DTT, 10% glycerol, 1 μg bovine serum albumin and 1.5 μg poly[d(I-C)] at room temperature for 30 min. Binding reactions were resolved in a 4% non-denaturing polyacrylamide gel (300 V, 90 min, 4 °C) in 0.5 × TBE. Gel was dried and exposed to X-ray film at -80 °C.

### Supplementary Figures

#### Supplementary Figure 1

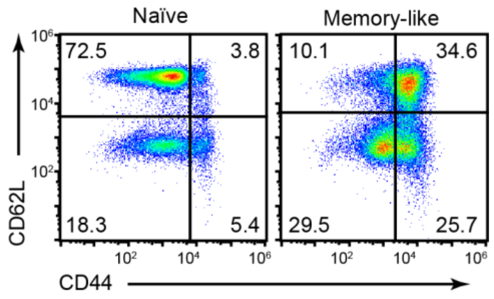


**Supplementary Figure 1.** Flow cytometry analysis of CD62L and CD44 surface staining showing naïve and differentiated memory-like $CD4^+$ T cell phenotype. Dot-plots are representative of three experiments performed.

#### Supplementary Figure 2

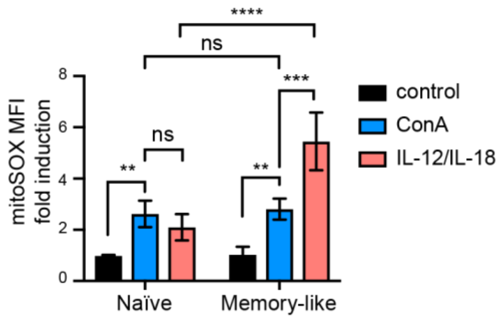


**Supplementary Figure 2.** Flow cytometry analysis of mitoSOX red fluorescence showing mitoSOX MFI fold induction (over unstimulated cells) in naïve and memory-like cells at 1 hour after ConA or Il-12 and IL-18 stimulation. The graphs show mean ± SD ($n$ = 3), *$p$<0.05, **$p$<0.01, ***$p$<0.001, ****$p$<0.0001, 2-way ANOVA (with Sidak's correction for multiple comparison).

#### Supplementary Figure 3

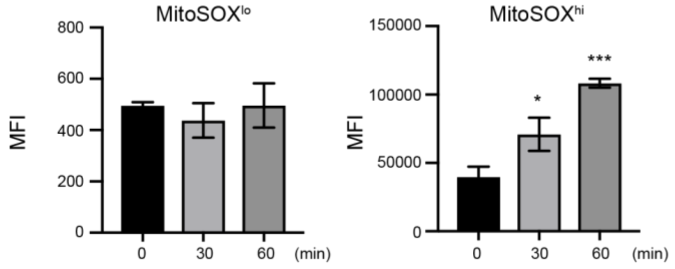


**Supplementary Figure 3.** Flow cytometry analysis of mitoSOX red fluorescence showing the MFI within mitoSOX$^{lo}$ (left) and mitoSOX$^{hi}$ (right) populations at early time points after IL-12 plus IL-18 activation. Graphs show mean ± SD ($n$=3); *$p$<0.05; ***$p$<0.001, one-way ANOVA with post-hoc Tukey test.

#### Supplementary Figure 4

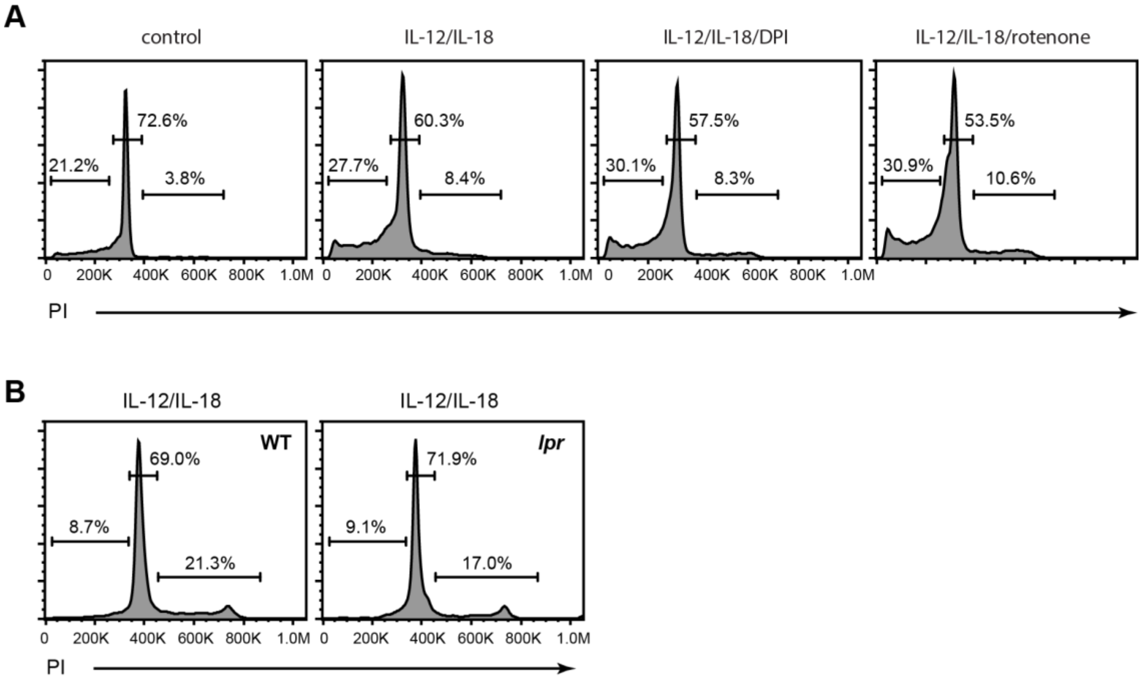


**Supplementary Figure 4.** Flow cytometry analysis of cell cycle after treatment with different inhibitors in WT cells **(A)** and in WT compared to *lpr* cells after IL-12 plus IL-18 stimulation **(B)**. Shown are representative data of 2 experiments performed.
